# Intestinal NF-κB and STAT signalling is important for uptake and clearance in a *Drosophila-Herpetomonas* interaction model

**DOI:** 10.1101/352369

**Authors:** Lihui Wang, Megan A. Sloan, Petros Ligoxygakis

## Abstract

Dipteran insects transmit diseases to humans, often in the form of trypanosomatid parasites. To accelerate research in more difficult contexts of dipteran-parasite relationships, we studied the interaction of the model dipteran *Drosophila melanogaster* and its natural trypanosomatid *Herpetomonas muscarum*. Parasite infection reduced fecundity but not lifespan in NF-κB/Relish-deficient flies. Gene expression analysis implicated the two NF-κB pathways Toll and Imd as well as STAT signalling. Tissue specific knockdown of key components of these pathways in enterocytes (ECs) and intestinal progenitor cells influenced initial numbers, infection dynamics and time of clearance. *Herpetomonas* triggered STAT activation and proliferation of Intestinal Stem Cells (ISCs). Loss of Relish suppressed the latter, resulting in increased parasite numbers and delayed clearance. Finally, loss of Toll signalling decreased EC numbers and enabled parasite persistence. This network of signalling may represent a general mechanism of the dipteran early response to trypanosomatids, crucial for parasite establishment and therefore transmission.

**AUTHOR SUMMARY:** Neglected Tropical Diseases are the most common diseases of the world’s poorest people. Many are caused by parasites called trypanosomatids that are transmitted to humans via insects belonging to the order of Diptera (also known as true flies). These flies (including tsetse, sand flies and black flies) are difficult to study in the lab and so the prospect of rapid progress in the basic biology of fly-parasite interaction is bleak. However, a model dipteran species with an extensive “tool-box” is the fruit fly *Drosophila melanogaster* with its natural trypanosomatid *Herpetomonas muscarum*. Here we establish the framework of their early interaction with the view that part of this interaction will represent an evolutionary conserved component of the dipteran response to parasite infection and will inform more targeted studies into medically important but difficult to study Diptera.

## INTRODUCTION

Neglected Tropical Diseases (NTDs) like sleeping sickness, leishmaniasis, hookworm infections, river blindness and elephantiasis are the most common infections of the world’s 1.4 billion poorest people and the leading causes of chronic disability and poverty [1,2]. NTDs are found mostly (but not only) in low and middle income countries [3]. For NTDs communicated to humans through an insect vector, the ability of the pathogen to overcome the insect’s midgut defenses is absolutely central to transmission. This is clearly illustrated by the following examples.

African trypanosomes, responsible for sleeping sickness and nagana, encounter a severe barrier to their establishment in the midgut of their tsetse fly vectors [reviewed 4]. It has been shown that there is increased resistance to *Trypanosoma brucei spp* (*T. brucei spp*) from their first blood meal where 50% of *T. brucei spp* become established, to their third blood meal onwards (the fly may take 40-60 blood meals in its life) where less than 10% of challenged flies become infected [5]. Paradoxically therefore, given their importance as vectors, tsetse fly populations are overwhelmingly resistant to trypanosome infection and the resistance mechanisms are manifested largely in the fly midgut [reviewed in 6]. However, our understanding of the molecular events underpinning this midgut-mediated resistance is poor [6,7].

Leishmania parasites seem to have successfully overcome barriers to establishment in their sand fly hosts as they develop in large numbers in the midgut of challenged laboratory strains. However, we understand neither how the parasite establishes itself in the gut nor why the insect tolerates large numbers of parasites. Nevertheless, in the wild there is only 1% of caught sandflies infected with Leishmania and there has been roles attributed to attachment of the parasite in the gut through lipophosphoglycan as well as insect galectins [reviewed in 8]. In the case of filariasis, the numbers of microfilaria ingested by all vectors (black flies, mosquitoes etc.) decline dramatically in the midgut lumen with either none or only a small fraction managing to penetrate the midgut barrier. Again, permissiveness of the mosquito midgut for parasite invasion is a key factor in determining success of the infections [9] but we understand virtually nothing at the molecular level of the mechanisms involved.

Unlike mosquitoes where technological development has been rapid, for some dipteran vectors challenged with kinetoplastid parasites the “tool box” required to tease out these interactions is very unlikely to be rapidly developed. For example, there is no realistic prospect of producing transgenic technology for tsetse flies because eggs are inaccessible due to intrauterine development of larvae; there is currently no transgenic technology for sandflies; maintenance of multiple lines of both flies permitting genetic studies is costly and complex; bioinformatics resources are in their infancy. In this context, the model dipteran insect *D. melanogaster* may be able to give answers on the possible existence of an evolutionary conserved component of the dipteran host response to kinetoplastid parasites.

Like all insects, *Drosophila* possesses a sophisticated antimicrobial defense. This is rapidly activated upon immune challenge by NF-*κ*B-like transcription factors through two distinct signaling cascades, namely the Toll and IMD pathways [reviewed in 10]. Sensing of *β*-1,3-glucan of fungi and peptidoglycan of Gram-positive bacteria principally trigger Toll signaling. This activation centers on the transmembrane receptor Toll, which is activated by the endogenous ligand Spz, a Nerve Growth Factor homologue [11]. Signal transduction through a receptor-proximal complex including Myd88, Tube and Pelle culminates in the proteolysis of the *Drosophila* IκB Cactus, which enables the translocation to the nucleus of the NF-κB homologue DIF [12]. Moreover, peptidoglycan from Gram-negative bacteria and Gram-positive bacilli primarily induce the IMD pathway, homologous to the TNFR1 pathway. Upon recognition by Peptidoglycan Recognition Proteins PGRP-LC and PGRP-LE, the signal is transmitted to Imd itself (a RIP-1 homologue) and then to TGF-β Activating Kinase 1 (TAK1), which activates the fly IκB Kinase (IKK) complex, which phospholylates the composite NF-κB/IκB Relish transcription factor [13]. The caspase-8 homologue Dredd cleaves Relish and releases the N-terminal DNA-binding part of the protein to translocate to the nucleus [14] and regulate hundreds of genes including several antimicrobial peptides (AMPs) [15]. In addition, the *Drosophila Ja*nus *K*inase/*S*ignal *T*ransducer and *A*ctivator of *T*ranscription (JAK-STAT) pathway has been implicated in defenses against viruses as well as a stress-response mechanism utilizing a set of core signaling components [reviewed in 16]. A transmembrane receptor encoded by *domeless* (*dome*), a single JAK tyrosine kinase encoded by *hopscotch* (*hop*), the transcription factor *stat92E* and *unpaired* (*upd*), as well as two related ligands encoded by *upd2* and *upd3*. Binding of Upd ligands to the Dome receptor leads to activation of Hop, which phosphorylates itself and Dome. Cytoplasmic Stat92E can bind to phosphorylated Dome/Hop complexes. Once bound to the Dome/Hop complexes, Stat molecules are phosphorylated and forming Stat dimers that will translocate to the nucleus and regulate target genes [16]. Both Imd and JAK/STAT pathway are involved in systemic as well as in epithelial immunity, especially gut epithelial immunity. There, the *Drosophila* midgut contains pluripotent intestinal stem cells (ISCs) that have a simple lineage: each ISC divides asymmetrically to produce itself and a transient enteroblast (EB), which will undergo terminal differentiation into either an polyploid absorptive enterocyte (EC) or as a diploid secretory enteroendocrine cell (EE) [17,18]. Individual ISCs are scattered along a thin layer of basal lamina in the posterior midgut and are the only proliferating cells in the epithelium [17,18]. This proliferation can be marked with an antibody against the phosphorylated form of Histone-3 (PH3^+^ cells) [19,20].

Very few studies of kinetoplastid interactions with *Drosophila* have been published. One biochemical study has looked at AMP production in response to infection with *Crithidia* spp [21]. In addition to not being a natural parasite for *Drosophila, Crithidia* largely infect the rectum of flies and are not a good model for midgut vector–parasite interactions. Nevertheless, natural gut-dwelling kinetoplastid parasites of *Drosophila* do exist [22]. A potential model system of greater relevance is *Herpetomonas ampelophilae*, a natural kinetoplastid parasite of *D. melanogaster*, which establishes infection in the midgut of the fly and can go on to invade the salivary gland [23, 24]. However, there are no studies that have examined the interaction between the adult *D. melanogaster* midgut and *Herpetomonas* beyond those initial papers. A recent study, has described the interaction between *Drosophila falleni* and *Jaenimonas drosophilidae*, a novel natural trypanosomatid parasite isolated from the wild [25]. There, *D. falleni* larvae were persistently infected throughout development, demonstrating persistent infection. However, there was a pronounced bottleneck in infection over metamorphosis with substantially lower rates of Infection in adults than in larvae (Hamilton *et al*, 2015). Using the rate of initial infectivity as a measure, these authors showed that *J. drosophilidae* infection in *D. melanogaster* larvae provoked an immune response that was not dependent on the IMD pathway [25].

In the present work we have developed a *Drosophila-Herpetomonas* system to be able to dissect insect-parasite interactions in a model dipteran insect. Using transcriptomics as well as tissue-specific RNAi assays, we have pinpointed the activation of the immune pathways responsible for gut defences upon parasite infection and documented the role of Toll, IMD and JAK-STAT signalling in modulating the number and time of parasite clearance. Linked to IMD-Relish, the timing of ISC proliferation had a pivotal role in the ability of the fly to clear the parasite quickly and to keep numbers down. Furthermore, our results provide a framework to establish the evolutionary conserved component of the response of dipteran insects against kinetoplastids.

## RESULTS AND DISCUSSION

**A natural parasite for *Drosophila melanogaster*** we have isolated trypanosomatid parasites from *D. melanogaster* caught in the wild on fruit baits in and around Oxford, UK (see materials and methods). Following sequencing of 18S rRNA we concluded that these parasites belonged all to the same species namely, *Herpetomonas megaseliae* (99% sequence identity with strain accession number U01014; data not shown) [26]. However, through the molecular redefinition of phylogenetic relationships in the *Herpetomonas* genus, *Herpetomonas megaseliae* is now included in *Herpetomonas muscarum* and henceforth we will call the isolated parasite *H. muscarum* [27]. In our hands, the prevalence of the infection in the wild was 5.01% in males (or 16 out of 212 male flies caught and identified as *D. melanogaster*), in accordance with reports of more extensive samplings [28]. The prevalence in females was approx. 6% (or 5 out of 91). Following species identification, males were assayed directly for the presence of the parasite and were used to isolate it. Female flies were kept to establish separate lines and verify insect species identification (see materials and methods).

We were able to culture the parasite (see materials and methods; Fig. 1A) and therefore wanted to transfer this host-parasite system in the lab to determine various aspects of this interaction. Pioneering work by Rowton and McGee has previously characterized infection of *Drosophila* laboratory populations by *Herpetomonas ampelophilae* [23, 24]. However, the experimental design did not preclude the infection of uninfected flies at different time points after the initial infection event. We therefore began by characterizing the likelihood, time course and fitness consequences of *H. muscarum* infection in laboratory *D. melanogaster* after a single exposure event. The infection protocol and scheme of time point sample collection for downstream experiments is illustrated in Fig. S1A. We also sequenced and annotated the genome of *H. muscarum*, determined its transcriptomic response inside the insect host and assayed its geographical distribution using publicly available sequencing data from *D. melanogaster* around the world (Sloan *et al.* in preparation).

**Figure 1.**
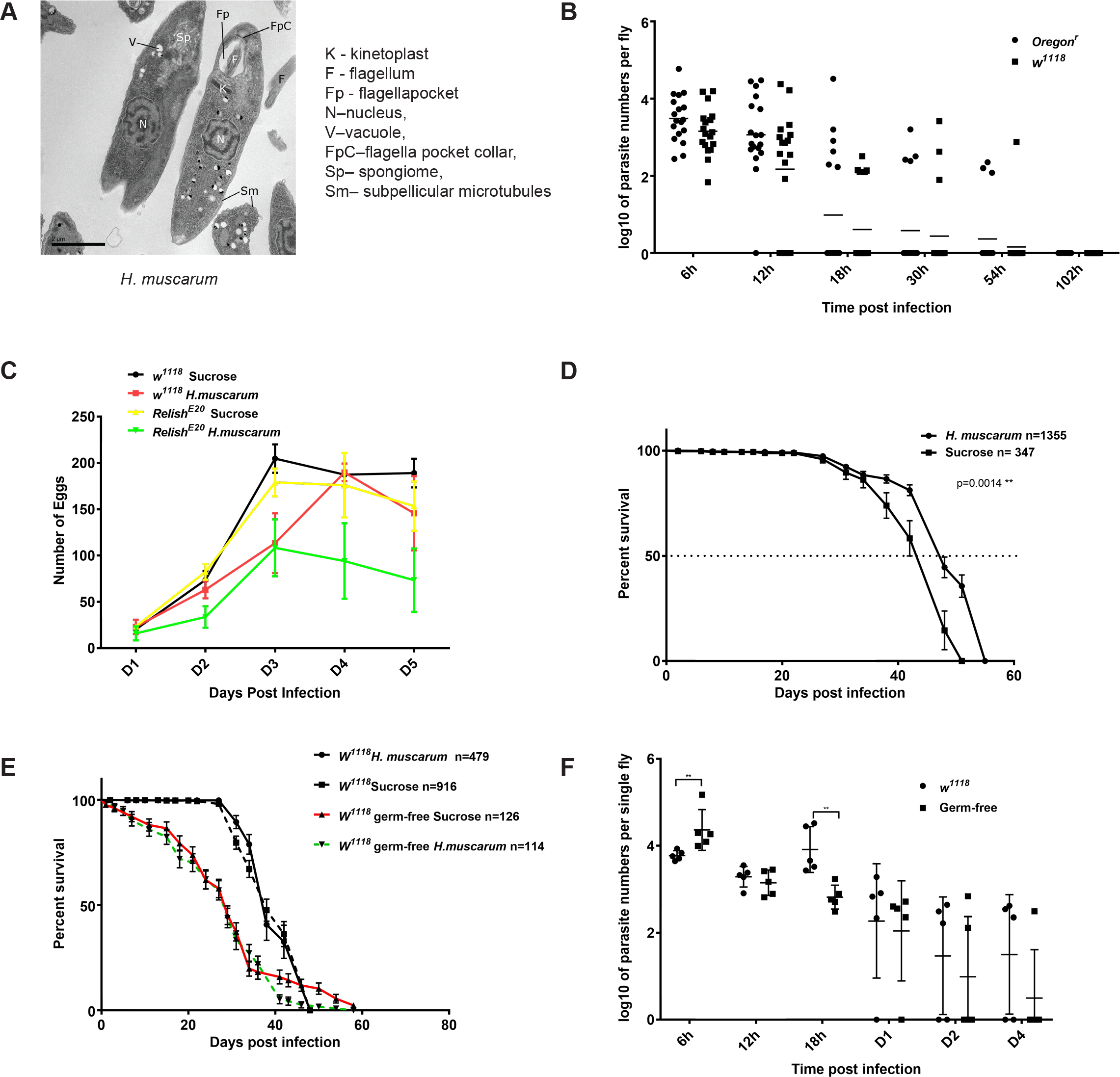
Infection of *Drosophila melanogaster* with its natural parasite *H.muscarum*. **(A)** EM of *H.muscarum* from culture. **(B)** Both *Oregon^R^* and *w^1118^* flies took up to 3 days to clear the parasites after an initial 6h oral feeding infection. **(C)** Fecundity assays of *w^1118^* and *w*^*1118*^; *relish* flies where infection reduced egg laying in both strains. **(D)** Life span of *Oregon^R^* was not affected after *H.muscarum* parasite oral infection compared to sucrose control. Infected flies lived significantly (but modestly) longer than controls. **(E)** Life span of *w^1118^* flies infected with *H.muscarum* was statistically indistinguishable compared to sucrose-only fed controls. Absence of gut microbiota showed a significant impact only late (LT_95_) in the lifespan of *w^1118^* germ-free flies compared to conventionally reared controls following *H. muscarum* oral infection. **(F)** In the absence of gut microbiota, more parasite intake was observed during the first 6 hours of feeding infection compared to conventionally reared flies. However, at 18h post-infection germ-free flies exhibited significantly reduced parasite numbers.

**Effects of *Herpetomonas* infection on *Drosophila* survival and life expectancy** We were able to record 100% infection rates in *D. melanogaster* laboratory flies fed with log-phase (72h) *H. muscarum* in 10% n “*i “i “i Q* sucrose (*Oregon^R^* and *w^1118^*; Fig. S1B). In this and all subsequent experiments, control flies are those fed with just 10% sucrose. Parasite presence was confirmed by visual inspection using live stains in the posterior (6h post-infection; Fig. S1C) and subsequently anterior midgut (24h post-infection) (Fig. S1D). These positions were reminiscent of the “swim back” of Leishmania towards the mouthparts in sand flies after digestion of the blood meal [8]. The possible attachment of *Herpetomonas* to the intestinal epithelium from the inside of the peritrophic matrix was also documented (Fig. S1E). Finally, the dynamics of infection was followed using quantitative real time PCR in reference to a standard curve that was made each time from a fresh parasite culture used for that specific infection experiment (Fig. S1F).

Infection dynamics showed that the majority of *Oregon*^*R*^ and *w*^*1118*^ flies were able to clear the parasite within 4-5 days although some flies were able to clear as fast as 12h (Fig. 1B). Parasite infection reduced egg deposition of *w^1118^* and *w^1118^; relish* mutant flies (Fig. 1C). This suggested a reduction in fecundity as a strategy to divert resources from egg production to parasite clearance [reviewed in 29]. Connected to the reduced energy spent for egg laying, *Oregon^R^* flies showed a statistically significant increase in their median lifespan when infected with the parasite compared to controls (Fig. 1D) while lifespan of infected *w^1118^* flies was statistically indistinguishable from controls (Fig. 1E). In germ-free conditions, *w^1118^* flies exhibited a higher initial parasite uptake (6h post infection) but later inhibited parasite proliferation, especially at 18h post-infection (Fig. 1F). Overall, there was a significant reduction in survival at the end of the lifespan curve (LT_95_; P<0.001) of parasite infected germ-free *w^1118^* compared to sucrose only fed controls (Fig. 1E).

Infected flies were not able to pass the parasite to non-infected adults when infected and non-infected flies were co-cultured in cages. To distinguish between infected and non-infected insects at the beginning of the experiment we marked the latter with an *Act-GFP* balancer chromosome (data not shown). This result showed that the parasite was able to survive inside one fruit fly host but could not subsequently transfer to an uninfected fly in contrast to *H. ampelophilae* [23]. In the latter case, transmission to adults and larvae was probably accomplished by feeding on substrates contaminated with host feces or “social digestion” of adult or larval cadavers by developing larvae. Our results however, leave open the possibility that flies killed all parasites prior to clearance or that an intermediate plant host is necessary for transmission, as some *Herpetomonas* species have been proposed to be associated with tomato plants [23]. Fast clearance and/or an intermediate plant host may also explain the low prevalence of infected fruit flies in the wild [30].

**Transcriptomic analysis of host response to parasite infection** We next sought to determine gene expression that was specifically altered by the presence of the parasite. We investigated transcriptome variations in whole, sucrose-only vs. sucrose-parasite fed flies following the same infection protocol shown in Fig. S1A. Transcriptome data were generated using the Illumina RNA sequencing platform of sequencing poly-A RNA (thus avoiding bacterial contaminants) to capture both known and novel coding and noncoding RNA (see materials and methods). Taking into to account that the process of parasite clearance took about 102 hours (see Fig. 1B) we followed infection dynamics of 4-day old flies from early time points (6, 12, 18h post infection), to 54h (as a mid-point) and finally to day 7, a time point after parasite clearance had been achieved. There were 1,556 genes that were significantly regulated following parasite infection (P<0.05). From these, our analysis identified 155 genes whose expression varied by at least a 2log change relative to expression in sucrose-only fed flies (Fig. 2A). The statistical confidence P-value was established in each of the three biological repeats. During the lifetime of infection those 155 genes included 66 upregulated-only transcripts, 56 downregulated-only transcripts and 33 (or 20%) that were upregulated in certain time points while downregulated in others. In terms of gene expression, the majority of the transcripts were differentiated at the initial stages of the response (98 and 79 at 6h and 12h following infection) while as the response progressed there was less differential expression (39 transcripts at 54h and 13 at Day 7, see Fig. 2B).

**Figure 2.**
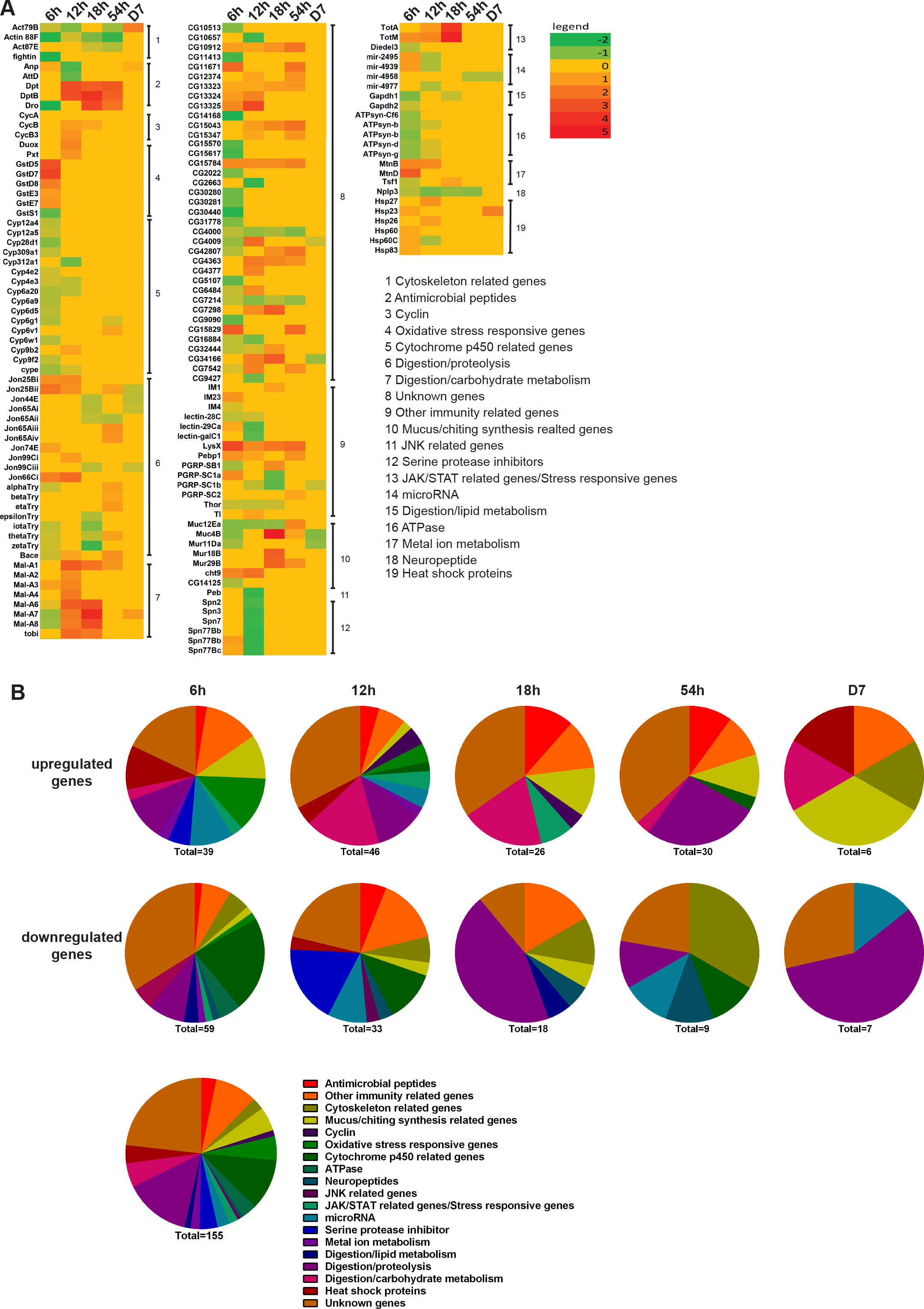
Host transcriptomics analysis of *H. muscarum* infection reveals the dynamics and gene networks involved. **(A)** Heat map showing the list of significantly regulated transcripts (>log2) compared to the corresponding sucrose control at each time point over the course of parasite infection. The list underlines the systemic temporal and spatial regulatory networks in response to *H.muscarum*. **(B)** This can be seen more clearly when the list in A is transformed into pie charts for every time point comparing percentage, gene numbers and gene ontology of transcripts both upregulated and downregulated over the time course of infection.

Using a global classification of gene ontology (GO), nearly a quarter (23.8%) of the genes were assigned as “unknown function” (11.6% upregulated-only, 8.38% downregulated-only and 3.82% both). Of those, the ones with at least one homologue could be divided into three main categories namely, intracellular signalling molecules, collagen-like cuticle proteins and genes with unknown function (our unpublished data). The rest of the transcripts were assigned to 17 functional categories (Fig. 2B). These included digestion related to proteolysis and lipid metabolism, oxidative stress responsive and serine protease inhibitors (all upregulated mostly at 6h post-infection), metal-ion metabolism (upregulated at 54h but suppressed at 18h and Day 7 post-infection), antimicrobial peptides (AMPs) and other known immune-related genes (including known pathways and stress responsive genes; gradually upregulated through to the mid-point at 54h post-infection), mucus/chitin synthesis related and cytoskeletal genes (upregulated at 54h and Day 7 but suppressed at earlier time points), neuropeptides (suppressed at all time points) and micro-RNAs (activated at 6h, 12h and 18h but suppressed at 54h and Day 7). Based on this analysis we concluded that parasite infection triggers high levels of signalling signatures associated to immune, stress and metabolic responses. In this work, we study the involvement of immune response genes.

**Immune pathways related to *Herpetomonas* infection in *Drosophila*** We found that following parasite infection, there were several differentially expressed genes including 1) AMP genes that were targets of the IMD pathway, 2) genes that coded for components of the oxidative stress response and 3) the Toll receptor itself. Therefore, we addressed the functional relevance of these findings using the GAL4-GAL80^ts^-UAS-RNAi system where we could knockdown genes in adulthood in a ubiquitous or tissue specific manner avoiding any developmental or long-term effects. At the permissive temperature (18°C) GAL80^ts^ prevents GAL4 binding and therefore RNA-interference, whereas shifting to the restrictive temperature (30°C) GAL80^ts^ releases GAL4 and thus induces RNAi [31]. We tested that the system was effectively induced using three GAL4 lines (see below): one expressed in all immunocompetent tissues (*J6-GAL4* or *C564-GAL4*) Fig. S2A), one expressed in enterocytes (*NP1-GAL4*; Fig. S2B) and one expressed in ISCs and EBs (*esg-GAL4;* Fig. S2C). The latter we also tested just at 18°C with infection vs. control to make sure that any difference we saw at 30°C was due to triggering RNAi and not a background effect. Indeed there was no expression of RFP in either infected or control guts at 18°C (Fig. S2D).

***Antimicrobials*** We found that silencing the gene coding for Dual Oxidase, the enzyme responsible for ROS synthesis in ECs [32], significantly increased initial loads (6h post infection) and delayed parasite clearance (Fig. 3A). Similarly, silencing *caudal* loss of which de-represses all Relish-regulated AMPs in ECs [33], significantly decreased parasite loads at early stages of infection (Fig. 3B). Conversely, silencing the AMPs *Diptericin* (Fig. 3C) or *Cecropin A1* (Fig. 3D) increased uptake and delayed clearance while overexpressing *Attacin* had the opposite effect (Fig. 3E). These results are consistent with a role for ROS as well as AMPs (especially those transcriptionally regulated by Imd/Relish) in initial uptake and clearing infection.

**Figure 3.**
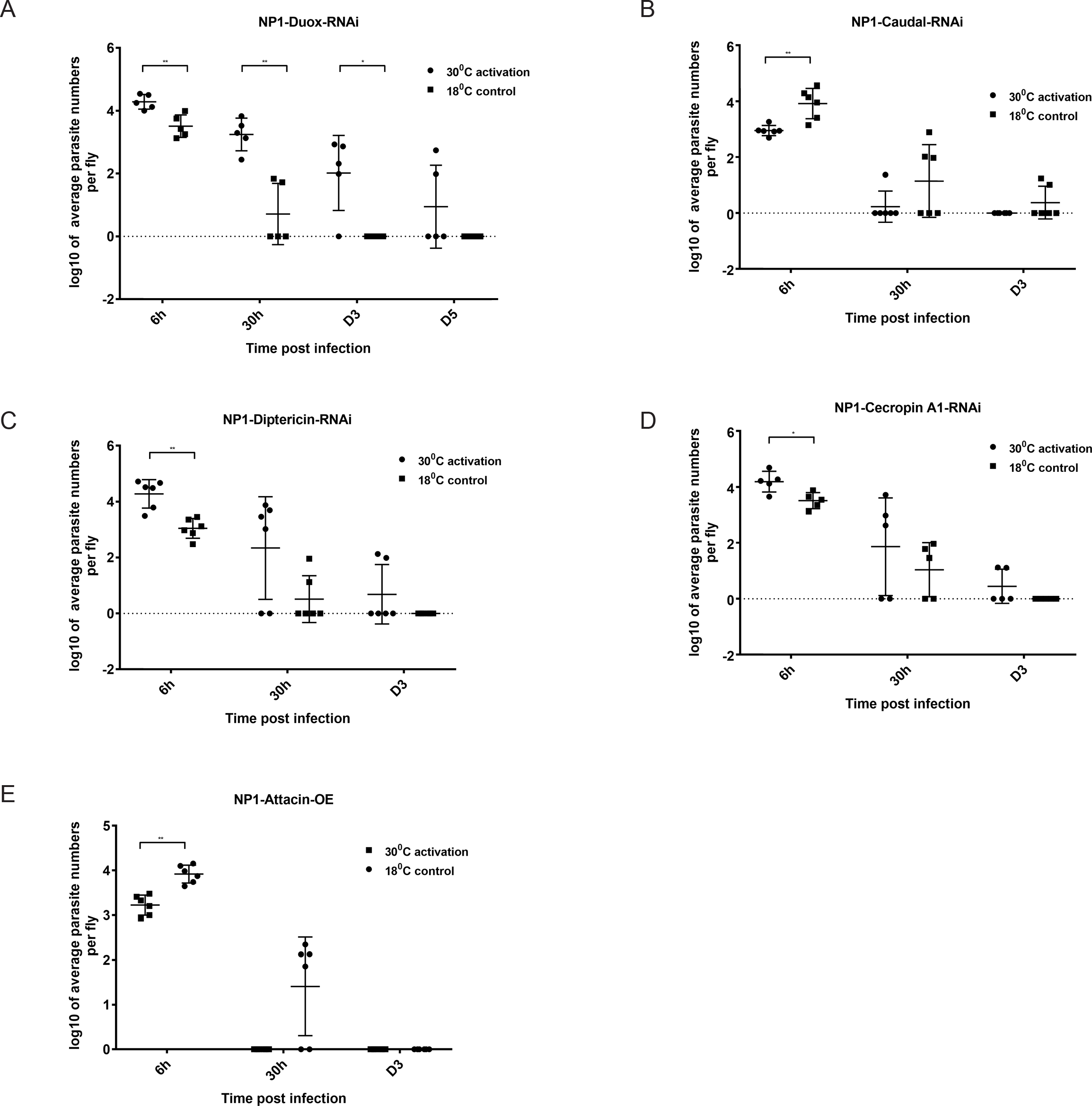
Influence of Dual Oxidase (Duox) and Antimicrobial peptides (AMPs) on *H. muscarum* intake, infection dynamics and clearance. **(A)** Activation of the GAL4/GAL80^ts^ system and silencing of Duox increased parasite numbers and delayed clearance beyond D5. **(B)** Derepressing transcription of a number of AMPs by knocking down *caudal* in ECs, reduced the parasite number intake and shortened the time for parasite clearance. RNAi in ECs (NP1-GAL4) of the AMP genes *Diptericin* **(C)** and *Cecropin A1* **(D)**, both targets of the IMD pathway, increased the initial parasite load. **(E)** In contrast, overexpression of the AMP gene *Attacin* in ECs cleared parasite infection in less than 30h.

***Relish*** Consistent with the results above, *Rel^E20^* flies displayed significantly higher parasite numbers at initial parasite intake at 6h compared to controls (Fig. 4A). In exploring the tissue-dependency for Relish in the control of parasite numbers, we found that silencing Relish in progenitor cells (ISCs and EBs) with *esg-GAL4* had a significant influence in parasite numbers throughout the lifetime of the infection and delayed parasite clearance by 48h (Fig. 4B). Conversely, overexpression in ISCs and EBs of a form of Relish that constitutively localises inside the nucleus [35] significantly decreased initial parasite loads and accelerated clearance with most flies clearing *Herpetomonas* at day 1 (Fig. 4C). In addition, silencing Relish in ECs significantly increased initial uptake (Fig. 4D) but no effect was observed when *rel* was silenced in the fat body (Fig. 4E). Therefore, Relish was necessary and sufficient to control parasite numbers in intestinal progenitor cells.

**Figure 4.**
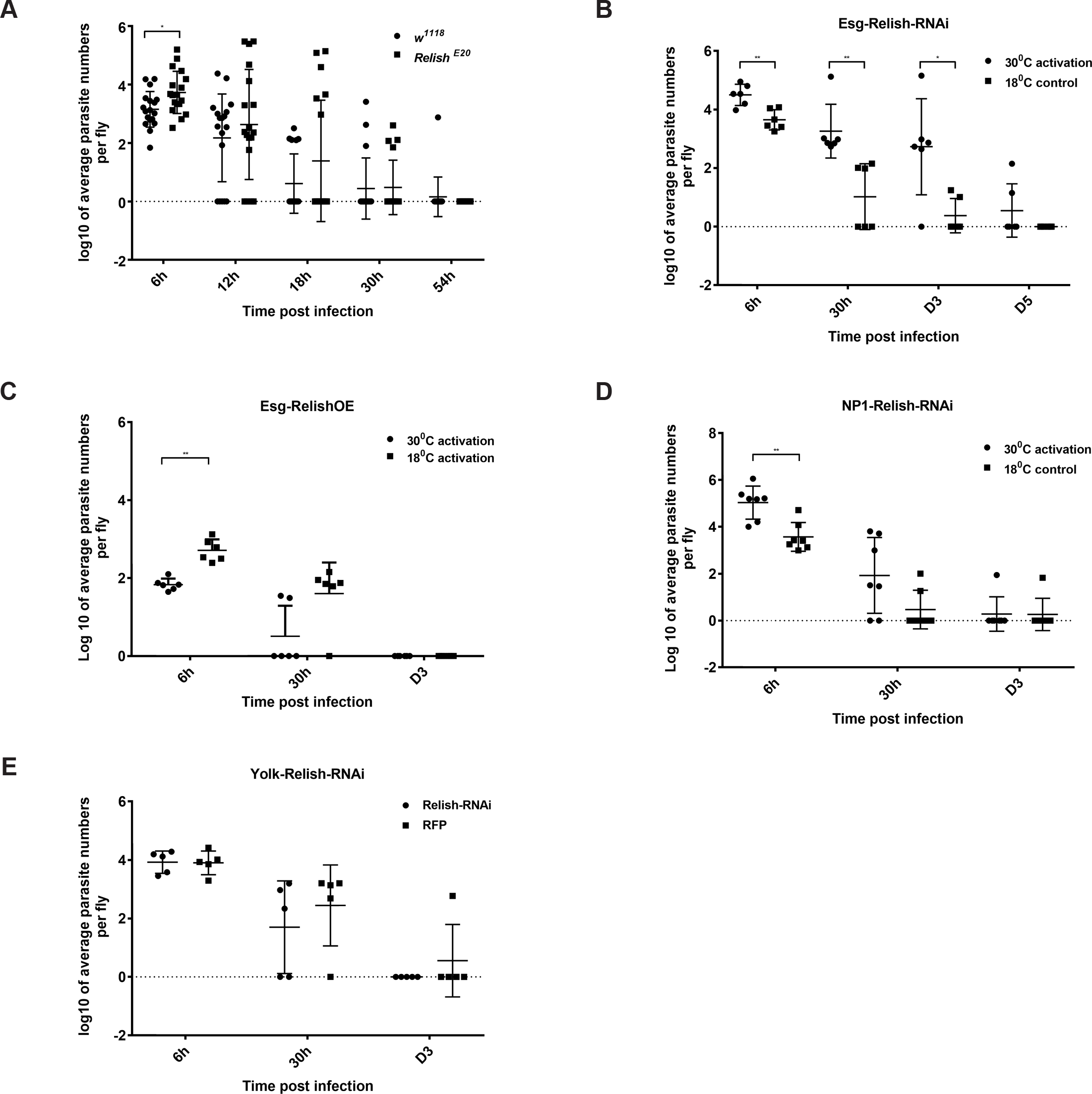
Influence of Relish on *H. muscarum* intake, infection dynamics and clearance. **(A)** Flies with loss of function of Relish (*Relish^E20^*), showed a significant increase in parasite number intake. Relish influenced the parasite intake and clearance in a tissue specific manner when silenced by RNAi: **(B)** a significant increase in both parasite numbers and clearance time was observed when Relish was blocked in intestinal progenitor cells. **(C)** Conversely, overexpression of Relish in progenitor cells (RelishOE) significantly decreased parasite intake. **(D)** A modest increase in parasite number intake was observed in the ECs. **(E)** No effect was observed in knockdown of Relish in immunocompetent fat body using the *yolk-GAL4* driver.

***Toll*** Next, we wanted to identify the role of the Toll receptor, which was differentially expressed in parasite-infected flies. Silencing Toll in all immunocompetent tissues significantly increased initial and early parasite load (6h to 1 day post-infection) (Fig. 5A). In contrast to Relish however, we found that Toll was not required in progenitor cells (Fig. 5B) but in ECs (Fig. 5C). In addition, silencing Toll expression in the fat body (using *yolk*-GAL4) indicated that Toll activity was also important there but only for controlling initial parasite load (Fig. 5D). To corroborate the role of the Toll pathway in controlling parasite numbers, we tested *dif^1^*, a mutant in DIF [11]. We found that *dif^1^* mutant flies displayed a significant increase in the initial *Herpetomonas* uptake (Fig. 5E). Moreover, *dif^1^* flies displayed a persistent infection that failed to clear at day 3 (Fig. 5E).

**Figure 5.**
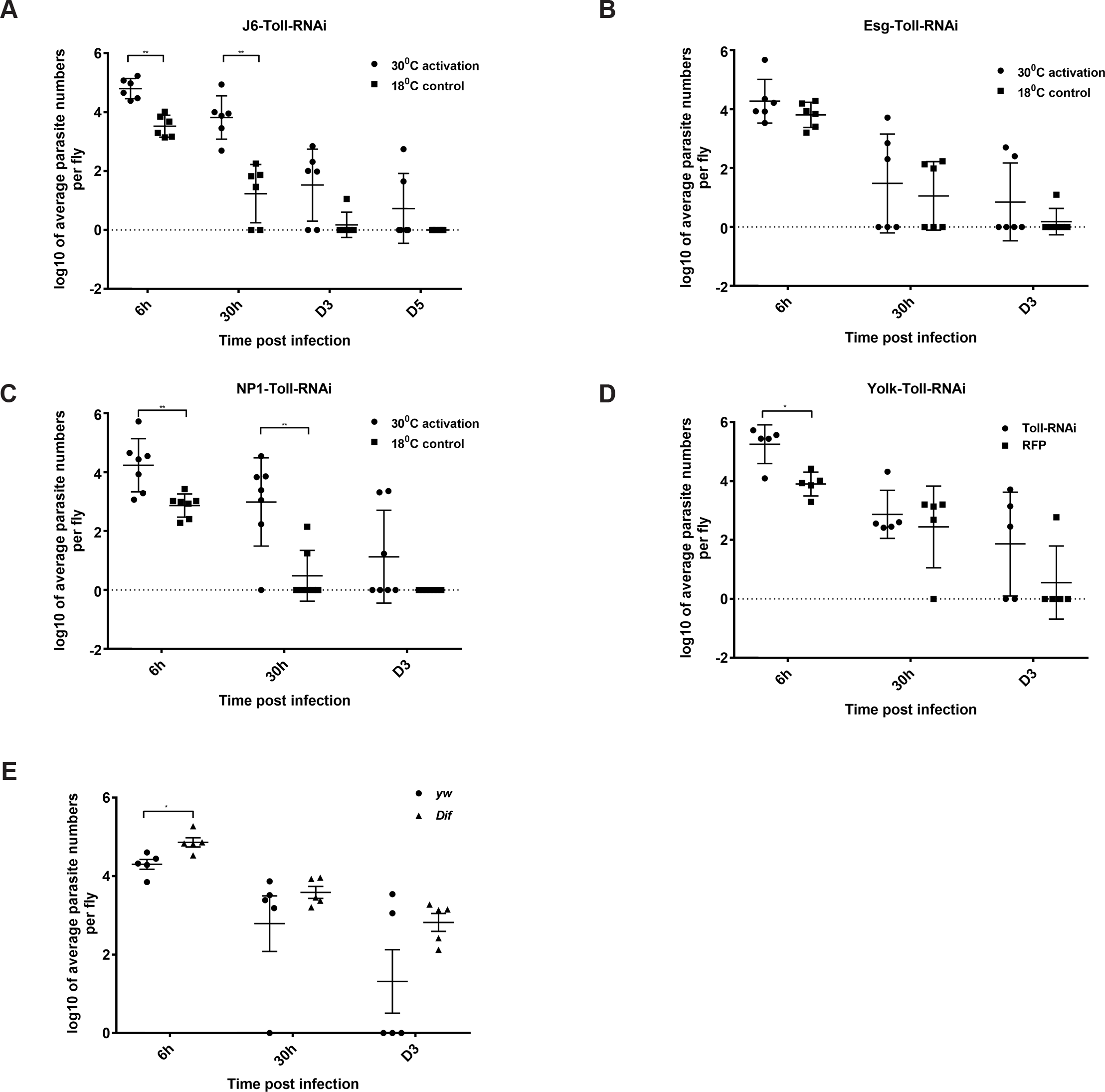
Influence of Toll on *H. muscarum* intake, infection dynamics and clearance. **(A)** Ubiquitous knocking down of Toll significantly increased parasite intake and slowed down clearance using the *J6-GAL4* driver. **(B)** Knocking down of Toll in progenitor cells did not significantly influence parasite intake or clearance time. **(C)** Knocking down of Toll in ECs increased the parasite intake and clearance time. **(D)** Absence of Toll in the fat body showed significant increase in parasite intake and slowed down the clearance compared to the RFP control. **(E)** Downstream of Toll, *dif* flies exhibited a significantly increased parasite intake and showed a persistent infection.

***STAT*** Targets and signaling components of the JAK-STAT were also differentially regulated following parasite infection. As the pathway has been implicated in gut physiology and immunity [16], we investigated the effect of silencing STAT in ECs, progenitor cells, hemocytes and fat body. Using J6-GAL4 we found that silencing expression of the Stat transcription factor (Stat92E) increased initial parasite numbers (6h post infection) and delayed clearance at 30h post infection (Fig. 6A). However, there was no requirement for STAT in ECs (Fig. 6B) or the fat body (Fig. 6C) or indeed the hemocytes (data not shown). However, silencing Stat in progenitor cells resulted in a significant increase in parasite numbers in the early phase of infection (6h-30h post infection; Fig. 6D). This result, along with the fact that Stat has been shown to be expressed in ISCs (see below) indicated that Stat activity was required in proliferating ISCs to control parasite numbers and clearance.

**Figure 6.**
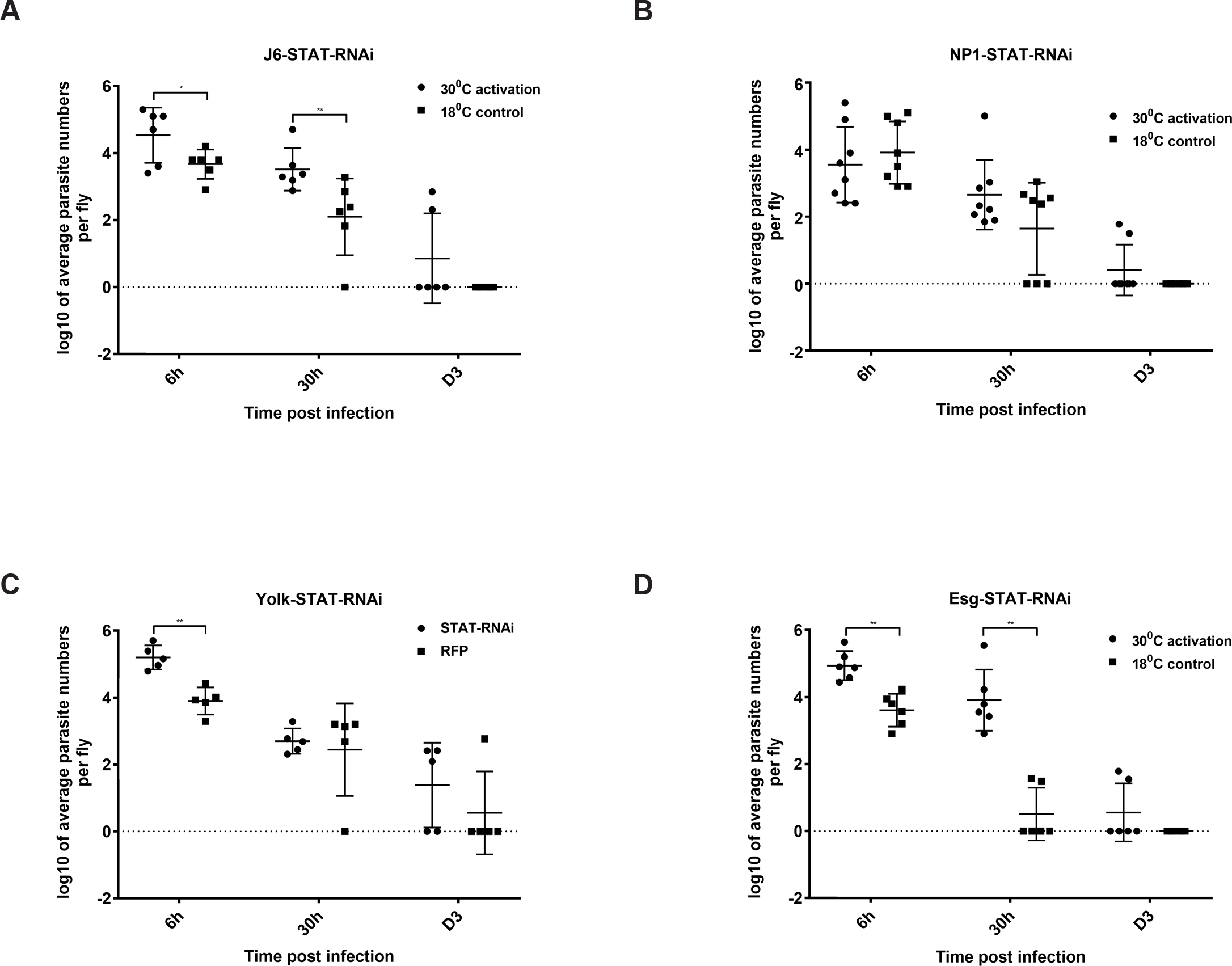
Influence of STAT transcription on *H.muscarum* intake and clearance. **(A)** Knocking down of STAT transcription factor in all immunocompetent tissues increased parasite intake and slowed down clearance. **(B)** Knocking down of STAT in ECs did not influence parasite intake number or clearance time. **(C)** When STAT was knocked down in the fat body where only parasite numbers at the earliest time point of 6h were increased. **(D)** The most prominent effect was observed when STAT was silenced in intestinal progenitor cells, significantly increasing both parasite number intake and clearance time.

**Parasite infection increases Stat-mediated transcription and proliferation of ISCs.** The involvement of JAK-STAT signalling in ISC proliferation and intestinal regeneration upon Upd cytokine secretion from ECs, has been documented previously [19, 34]. We documented the involvement of JAK-STAYT signalling during parasite infection by examining a reporter of JAK-STAT signalling activity, where multimerized Stat92E consensus binding sites control expression of destabilised-GFP (lOx-STAT-GFP) [36]. Following parasite feeding, GFP-expressing cells were increased during the early phase of infection, with significant difference from the control at 6h and day-3 post infection (Fig. 7A). This correlated with a significant increase in proliferative ISCs measured with an antibody against phospho-histone-3 (PH3) (Fig. 7B). To verify that the PH3 result was not a genetic background effect, the statistically significant increase in ISC proliferation was also verified using PH3 in infected vs. sucrose-only fed *w^1118^* flies (Fig. 7C). Moreover, there was a significant increase in GFP-positive progenitor cells (ISCs and EBs) when an *esg-GAL4; UAS-GFP* strain was infected with *Herpetomonas* (Fig. 7D).

**Figure 7.**
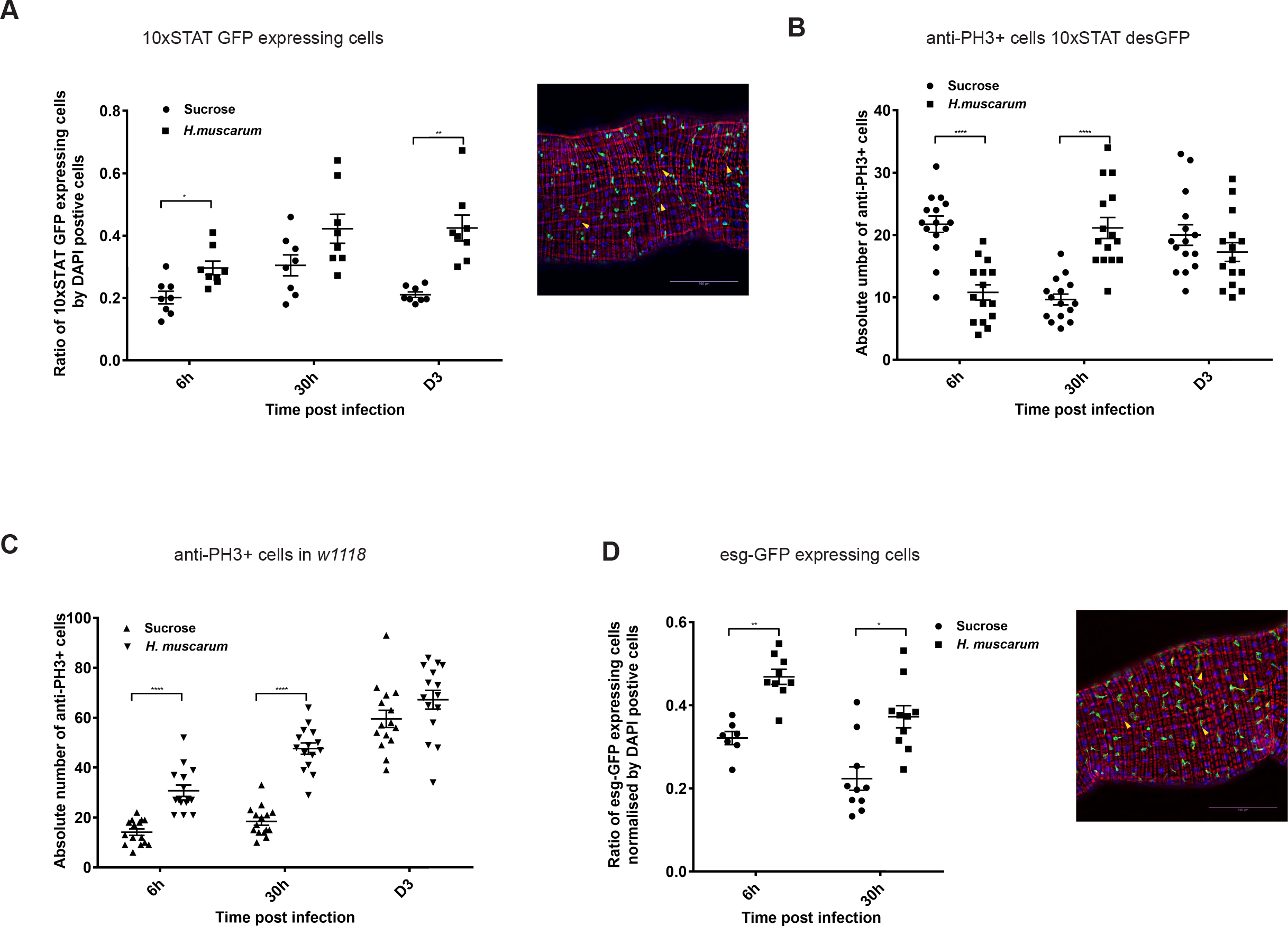
STAT-mediated transcriptional activation & ISC proliferation following *H.muscarum* infection. **(A)** Significantly increased JAK-STAT activity was quantified by measuring the GFP expressing cells in the gut following parasite infection in a reporter line where GFP was under the control of 10 copies of STAT binding sites (*10xSTAT-desGFP*). A selected region of the midgut at 20x magnification is shown (right panel). Yellow arrows indicate the GFP cells co-localizing with DAPI which were counted. Quantification of GFP expressing cells normalized with DAPI from 8 guts is shown (left panel). Results in the graph are from three independent infections. Following parasite infection, the number of GFP-expressing cells showed significant upregulation at 6h and D3 post infection. **(B)** The number of proliferating ISCs identified by anti-histone 3 (anti-PH3) antibody in infected *10xSTAT-desGFP*, showed significant increase at 6h and D1 post infection. **(C)** With the same technique as in B, increase of proliferating ISCs could be quantified in *w^1118^* flies. **(D)** Increased number of ISCs and EBs following parasite infection at day 1, was quantified by measuring GFP expressing cells in the gut of *esg*-CD8-GFP, *UAS-Gal4*, *gal80*^*ts*^ flies after an initial 6-day incubation at 30°C (infection and subsequent culture at 25°C). A selected region of the midgut at 20x magnification is shown (right panel). Yellow arrows indicate counted GFP cells co-localizing with DAPI. Quantification of GFP expressing cells normalized by DAPI from 8-10 guts and three independent infections is shown in the graph (left panel).

**Loss of Relish suppresses ISC proliferation following parasite infection** We next explored the signaling underlying proliferation of ISCs following parasite infection. When *w^1118^* were infected with *H. muscarum*, PH3^+^ cells were significantly increased continuously compared to the sucrose control, starting from 6h (Fig. 8A), day 1 (Fig. 8B) and day 3 (Fig. 8C). In contrast, sucrose-only fed controls showed a stable number of PH3^+^cells in both 6h (Fig. 8A) and day 1 (Fig. 8B) increasing only at day 3 (Fig. 8C). The number of PH3^+^ cells in sucrose-only fed *imd^R156^* was significantly higher than *w^1118^* controls indicating a de-repression of progenitor cell proliferation in the absence of Imd (Fig. 8A). Parasite infection significantly suppressed ISC proliferation in the absence of Imd at 6h post-infection, indicating a strategy from the parasite’s side to achieve midgut establishment (Fig. 8A). Nevertheless, this suppression was not evident at day 1 (Fig. 8B) or day 3 (Fig. 8C) presumably because the *imd* allele used was a hypomorph. At these time-points, the numbers of PH3^+^ cells between infected and sucrose-fed *imd^R156^* flies were statistically indistinguishable. Intestinal proliferation in *rel^E20^* mutant flies (sucrose control) was also significantly increased compared to *w^1118^* sucrose controls but not as pronounced as *imd*^R156^sucrose controls (Fig. 8A). As with *imd^R156^*, loss of Relish significantly suppressed ISC proliferation following parasite infection (Fig. 8A). The number of PH3^+^ cells in *rel^E20^*-infected intestines was kept significantly lower both at day 1 (Fig. 8B) and day 3 (Fig. 8C) compared to infected *imd*^R156^ and controls. When silencing *rel* in progenitor cells, the majority of the GFP expressing cells in the guts of esg-CD8-GAL4, UAS-GFP; GAL80^ts^ flies were enlarged GFP-positive cells that had lost their proliferative capacity as no PH3^+^ was detected (Fig 8D). Conversely, overexpression of Relish increased the population of smaller GFP expressing ISC-looking cells, which correlated with an increase in PH3^+^ cells in sucrose-fed controls (Fig. 8E). However, this PH3^+^ proliferation was suppressed by the parasite (Fig. 8E).

**Figure 8.**
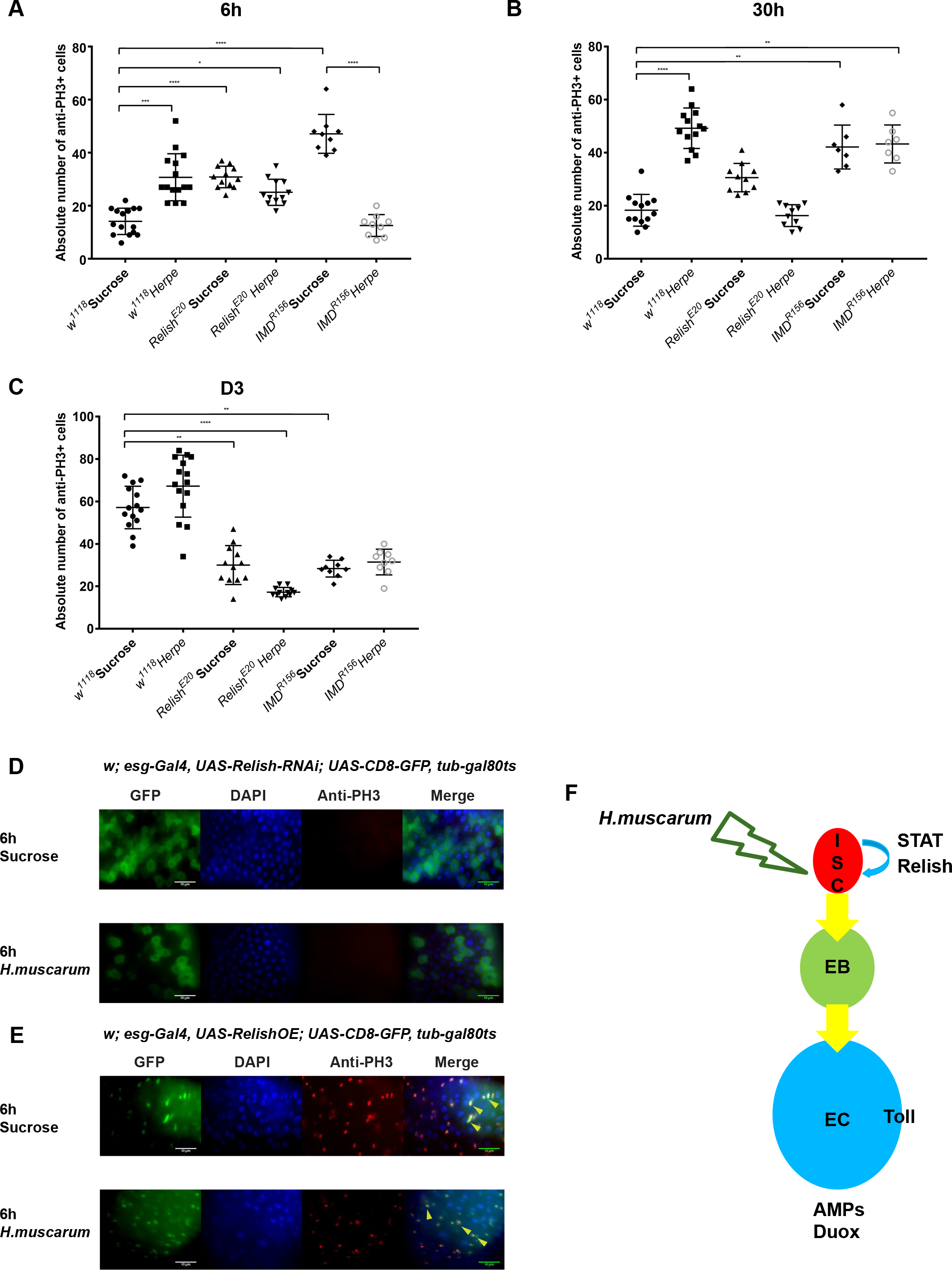
ISC proliferation and parasite infection. ISC proliferation dynamics following *H.muscarum* infection at 6h **(A)**, Day 1 **(B)** and Day 3 **(C).** ISC proliferation was induced following parasite infection but blocked when Relish was knocked-down. **(D)** The effect of Relish-RNAi was confirmed at the level of the tissue (lOOx magnification of anterior midgut region) where loss of Relish abolished anti-PH3 staining (red) and increased large non-proliferative EB-like esg-GFP-positive cells. **(E)** Conversely, overexpression of Relish induced proliferation (red) and increased small ISC-like *esg*-GFP-positive cells. Yellow arrows show the co-localization of PH3 and GFP staining confirming ISC status. **(F)** A cartoon illustrating a working model based on the current finding on the *H. muscarum* oral infection model in the *Drosophila* gut. Both JAK-STAT signaling and Relish transcription factors are essential in ISC proliferation. The combinatorial effect from JAK-STAT signaling and Relish transcription factor encourages a hyper proliferation of ISCs, which eventually results in a fast epithelium turn over in the *Drosophila* gut to remove the parasite. Together with the expression of effector molecules such as AMPs and the peristaltic movement of the gut the parasite can be cleared quickly and efficiently.

Thus, independently of infection, Relish was sufficient for ISC proliferation. High levels of Relish in the nucleus pushed cells to be PH3^+^, with ISC morphology whereas low levels of Relish provoked progression in the progenitor lineage to larger EBs (*esg*-positive non proliferative cells). The role of Relish in proliferation of midgut progenitor cells and the fact that its loss resulted in sustained low numbers of PH3^+^ cells following infection, suggested an explanation for the delay in parasite clearance in *rel^E20^*. In addition, STAT-mediated transcriptional activity correlated with ISC proliferation following infection and was important for parasite clearance. Finally, AMPs, Duox/ROS and Toll were required in ECs (Figure 8F).

Nevertheless, even in the absence of Relish the parasite was eventually cleared. This suggested that gut epithelium renewal was just one of the factors influencing parasite clearance and that most likely “flushing” of the parasite through the mechanical contraction of the gut was at play. Clearance by “flushing” would be possible if the parasite was failing to attach to the gut epithelium. Nevertheless, there is significant upregulation of *Herpetomonas* genes inside the host that are homologous to attachment genes in other trypanosomatids (Sloan *et al*, in preparation). More work is needed to define a possible interaction interface for this attachment.

*Herpetomonas* infection induced an acute early increase in ISC proliferation in *w^1118^* control flies as seen with PH3^+^ quantification (Fig. 8A). However, this was suppressed in immune-deficient *rel^E20^* or *imd^R156^* flies even though these flies displayed a higher than the control ISC proliferation in sucrose-only treatment (Fig. 8A). This suggested that the parasite actively suppressed proliferation of ISCs and epithelial renewal as a means of establishing a stable midgut presence and the Imd/Relish pathway was paramount to resist this strategy. This has been also observed in *Vibrio cholera* [37] and *Pseudomonas entomophila* [38] intestinal infections. It would be interesting to see whether suppression of ISC proliferation happens in the intestines of tsetse infected with trypanosomes or sand flies infected with *Leishmania*.

**Conclusions** We established for the first time a framework to systematically study how the model dipteran insect *Drosophila*, responds to a trypanosomatid parasite infection. We showed that Relish, STAT, Toll, Duox and AMPs were all important for fast clearance and control of parasite numbers. Our results provide cellular context to similar data from tsetse where RNAi of Relish or of an AMP homologous to *attacin* increased *T. brucei* numbers in infected tsetse flies [39]. Thus, the *Drosophila-Herpetomonas* system could be used to address gene function questions in insect vectors (e.g. tsetse, sand flies, horse flies, black flies etc.) where the tool-box available for direct functional studies is not yet fully developed while the establishment and maintenance of insect colonies is difficult.

## Materials and Methods

***Drosophila* collection and species identification** We collected *Drosophila* in a residential area using the corresponding author’s back garden and compost tip (inside the garden) as places to trap fruit flies. As such, no consent was necessary. Collection was done through 5-day periods from late March to late June 2011. During this period, traps containing fermented banana were set every week. The nine species of the *melanogaster* species subgroup are morphologically very similar but male genitalia is a reliable characteristic to distinguish males [28]. For females we used as a discriminatory characteristic the area below the eye (cheek) along the long axis, which is broader in *D. melanogaster* than all other members of the group [40]. To confirm our identification following establishment of separate stocks from a single female founder we selected flies randomly identified morphologically and used a PCR diagnostic test for the antimicrobial peptide gene drosomycin as described [22]. Species identification was confirmed with first generation offspring.

***Drosophila* stocks** Single trapped females were isolated in vials and their offspring cultured as separate “isofemale” lines to verify species identification [28]. Trapped males were used to isolate and culture the parasite. Starting from a single cross (one female-one male), *Oregon^R^* flies were used as a wild type laboratory strain, for establishment of laboratory oral feeding infection protocols and transcriptomics experiments thus ensuring a less variable, streamlined genetic background. *w^1118^* {BL #6326) and *yw^67c23^* (BL #6599), were used as controls and for the genetic background of all the other strains used in these studies and were obtained from the Bloomington Stock Centre. IMD, Toll and STAT signalling pathway mutants: *yw^67c23^*,*Dredd*^*B118*^ (BL #55712); *w^1118^*;*relish*^*E20*^ (BL #55714), *y*^*1*^ *w*\*; *Dif*^*1*^ *cn*^*1*^ *bw*^*1*^ (BL #36559) and *y*^*1*^ *v*^*1*^ *hop*^*Tum*^ (BL #8492) were also obtained from the Bloomington Stock Centre. The GAL4 driver lines used were: *w*^*1118*^, *np1-GAL4* (ECs) was kindly provided by Heinrich Jasper (Buck Institute for Research on Aging, Novato, CA, USA), *yolk-GAL4* (female fat body), *w*^*1118*^; *P{GawB}c564* (*J6-GAL4*, expressed in fat body, gut, hemocytes, (BL#6982) and *esg-GAL4* (expressed in ISCs and EBs) was obtained by Bruno Lemaitre EPFL, Lausanne see ref [19].

All the *w*^*1118*^, *UAS-RNAi* transgenic lines were purchased from Vienna *Drosophila* Resource Centre (VDRC). *GAL80^ts^* strains were all obtained from Bloomington Stock Centre: P*{tubP-GAL80ts}Sxl*^*9*^, *w*\*/*FM7c* (BL*7016), *w*\*; *P{tubP-GAL80tsf/TM2 (BL#7017)*, *w*\*; *sna*^*Sco*^/*CyO*; *P{tubP-GAL80ts}*^*7*^ (BL#7018) and w*; P{tubP-GAL80ts}^20^; TM2/TM6B, Tb^1^ (BL#7019). Appropriate lines were used to cross with the *GAL4* driver lines to ensure that the *GAL4* and *GAL80^ts^* genes were bred onto different chromosomes into homozygosity before being used in RNAi screening and other breeding schemes. We also used *10xSTAT-desGFP* [36].

**Culture of *H.muscarum*** The isolated *H.muscarum* used in the study was routinely cultured in 10% FBS (Gibco) supplemented *Drosophila* Schneider’s-2 (S2) media (Sigma-Aldrich). The parasite culture was maintained by sub-culturing every 3 days at 1:100 dilution to a fresh 5 ml S2 media. To count the parasite cells, cultures were first spun down at 2,000rpm for 5 mins and the cell pellets were treated with 10% methanol for 15 minutes to kill and immobilize the parasite cells. The cells were subsequently precipitated by spinning at 2,000rpm for 5 mins and washed twice by 1xPBS. After washing, cell pellets were resuspended in 1XPBS and diluted by 100 times in PBS before being transferred to a hemocytometer to count under a bright field light microscope at 100x magnification.

**TEM Microscopy of *H. muscarum* cells:** 5mls of culture at 5E6 cells/ml was fixed by adding 2mls of 16% PFA and 0.8 ml of 25% glutaraldehyde. This was incubated at 28°C. Cells were pelleted at 10000rpm for 5 mins and the fixative removed. Cell pellets were washed twice in 1ml 0.1M PIPES pH7.2 and once in lml 0.1M PIPES pH 7.2 with 50mM Glycine. The supernatant was removed and the pellets embedded in 0. 5ml 2.5% agarose. The pellets were trimmed down and fixed in OsO4 1% for 1 hour at 4°C, washed three times in 2ml milli-Q water and then incubated o/n in 0.5% Uranyl acetate in milli-Q water at 4°C. Cell pellets were washed for 10 mins in 2ml of milli-Q water at room temperature on a rotor before a 30min incubation in 30% EtOH on ice. The pellets were then dehydrated by immersion in increasing concentrations of 2ml EtOH (50%, 70%, 80%, 90% and 95%) for 10 mins each on ice. The three final dehydrations were done for 30min in 2ml 100% dry EtOH. Pellets were then incubated in 2ml of the following mixes of ethanol: agar100 resin; 3:1 (1 hour with rotation), 1:1 (2.5 hours with rotation), 1:3 (1 hour with rotation). Pellets were then incubated in 2ml 100% agar100 resin for 24 hours with the resin replaced with fresh resin at 18 hours and 22 hours. Finally pellets were transferred into the base of beem capsules and the resin set at 60°C for 18 hours. Images were taken using the FEI Technai 12 Transmission electron microscope with the Gatan Ultrascan 1000 CCD camera and Gatan Digital micrograph and SerialEM image acquisition platforms.

**Oral feeding infection and quantification of parasite by real-time PCR** For each independent infection of a group of 20-30 flies, 10e7 *H. muscarum* parasite cells were harvested from a 3 days-old culture (which showed the highest infectivity rate from our experience) and resuspended in 500ul 1% sucrose. The parasite solution was then transferred to a 21mm Whatman Grade GF/C glass microfibre filter circle (Fisher Scientific). Circles containing the parasite cells were placed into standard *Drosophila* small culture vial without any food. The flies used in the infections were 4-5 days old before they were starved overnight. After starvation, the flies were transferred to food vials that contained the Whatman circles with the parasite cells. After 6h of feeding, flies were moved and reared on standard yeast/molasses medium. At different time points post oral infection (including the 6h feeding as 6h point), infected flies were collected for downstream experiments.

An absolute quantification of real-time PCR based method was developed to quantify the parasite numbers in the flies. For each infection, *H.muscarum* parasite cells from 1ml of 3 days old culture were harvested and counted as described as above. The genomic DNA of the cells was extracted using cells and tissue genomic DNA isolation kit from Norgen Biotek following the manufacturer’s instruction. The purified gDNA was serially diluted and was used as templates in RT-PCR reactions to obtain linear regression of gDNA verse Ct values using *H.muscarum* gene specific primers. The serial dilution of parasite gDNA was then correlated with the serial dilution of the absolute parasite cells. Hence a stand curve of the parasite numbers verse the Ct values can be established as shown (Fig. S1E). For each individual infected fly or a group of flies, gDNA from both the fly and the parasite was exacted with the cells and tissue genomic DNA isolation kit. By measuring the Ct value of the gDNA sample, the absolute number of parasite cells could be extrapolated from the stand curve. *H. muscarum* gene specific primers used in the RT-PCR was designed based on the paraflagellar rod protein (PFR2) gene (GenBank accession no: AY785780). The primer sequences were: HmegRodF 5’-GGACTGCTGGAACAAGATC-3’, HmegRodR 5’-AGCTTCTTGTGCTGGGAG-3’.

**RNAseq and data analysis**_Total RNA of 8-10 flies at 6h, 12h, 18h, 30h, 54h and D7 (Fig. S1A) and post *H. muscarum* oral infection was extracted with total RNA purification kit from Norgen Biotek following the manufacturer’s instruction. Triplicate of samples were used for each time point. cDNA libraries were prepared with the Illumina TruSeq RNA Sample Prep Kit v2. All sequencing was performed on Illumina HiSeq 2000 plaftform using TruSeq v3 chemistry (Oxford Gene Technology, OGT). All sequence was paired-end and performed over 100 cycles. Read files (Fastq) were generated from the sequencing platform via the manufacturerɹs proprietary software. Reads were processed through the Tuxedo suite [41]. Reads were mapped to their location to the appropriate Illumina iGenomes build using Bowtie version 2.02. Splice junctions were identified using Tophat, version v2.0.9. Cufflinks was used to perform transcript assembly, abundance estimation and differential expression and regulation for the samples (verstion 2.1.1). RNA-Seq alignment metrics were generated using Picard.

At each corresponding time point, total RNA from triplicate of sucrose fed controls was extracted and the transcriptomes were sequenced and analyzed in the same manner for the subsequent comparison and data analysis.

**Lifespan monitoring** Life span of flies infected or sucrose fed controls was monitored by counting the number of live flies on a 3-day interval until the last single fly in the culture died. During that time live flies were put in fresh food every 2 days.

**Generation of germ-free flies and RT-PCR to measure dynamics of gut microbiota** Germ free flies were generated following standard protocols [19]. Briefly, 200-300 embryos were collected from fly cages fitted with apple juice plates. The embryos were then treated with 50% bleach for 2-3 minutes until all embryos were dechrionated when observed under a stereoscope. Subsequently, embryos were washed twice with 70% ethanol followed with autoclaved mili-Q water to remove residual ethanol. Treated embryos were then transferred to vials with standard yeast/molasses medium. Both vials and food were autoclaved forehand. From then on, aseptic procedures were employed in handling of vials with flies.

**Dissection of fly guts and immune-staining** Dissection of the fly guts and immune staining was generally following a standard protocol as in [41]. Rabbit anti Phospho-Histone H3 (Ser10) (anti-PH3) against *Drosophila* and goat anti rabbit Alexa Fluor A568 secondary antibody was obtained from Invitrogen.

**Fluorescence microscope and image analysis** Images of the gut with GFP expressing cells, DAPI and anti-PH3 staining were obtained with either Zeiss Axioplan 2 (Carl Zeiss) or Zippy DeltaVision Elite (Applied Precision). Image J was used to quantify of individual cells from the stacks taken by Zippy DeltaVision Elite. GFP expression cells and DAPI cells in the same designated region were counted separately and automatically. The relatively ratio of GFP expressing cells normalized by DAPI cell was used instead of absolute cell numbers to represent more accurately the change of the GFP expressing cells by eliminating factors such as physiological and anatomical change of the gut tissue in response to parasite infection and/or experimental handling.

**Statistical Analysis** Data in this study were analyzed using GraphPad Prism version 7.00 for Windows (GraphPad Software, La Jolla California USA, www.graphpad.com). The mean values were compared by nonparametric unpaired Mann-Whitney test between two groups. In all tests, * indicates P ≤0.05 which was considered significant. ** indicates p≤0.01 *** indicates p≤ 0.001 and **** indicates p≤0.0001. In all the experiment settings, at least three independent repeats were performed throughout the study.

<u>Survival data</u> All life span data were analyzed using GraphPad Prism (version 7). Briefly, we pooled the lifespan data (n indicated in the figures) from at least 3 independent biological repeat. Sets of life span assays for each genotype or each experiment setting as in germ-free vs conventional reared flies (i.e. there were no significant departures between repeats; P>0.1, log-rank test) were compared. Kaplan Meier estimates were used to plot survival curves for each experiment. Significant differences between survival data of different genotypes with or without parasite infection were identified using Log Rank and Wilcoxon tests (ChiSquare and p-values).

<u>RT-PCR data</u> For RT-PCR analysis to quantify the average absolute number of parasites present in the flies, we performed 5 to 6 biologically independent measurements with n=3-6 for each independent experiments. For Fig 1B, RT-PCR analysis to quantify the absolute number of parasites present in single individual fly was performed. We pooled results from 3-4 independent experiments with the total n=18-22. All values were expressed as mean values and were plotted with standard error. Nonparametric unpaired Mann-Whitney test were employed to ascertain statistical significance.

<u>Gut image and anti-PH3+ cell analysis</u> To quantify and compare the GFP expressing and anti-PH3+ cells, we pooled and combined the guts from at least 3 independent experiments with the n=6-15. Cells were counted as described in the materials and the methods. In Fig. 7 A-D, GFP expressing and anti-PH3+ cells with or without parasite infection were compared at 6h, 30h and D3 post parasite challenge. Nonparametric unpaired Mann-Whitney test was employed to ascertain statistical significance. In Fig. 8 A-C anti-PH3+ cells were compared and analyzed at 6h, 30h and D3 post challenge. All values were expressed as mean values and were plotted with standard error. One way ordinary ANOVA were employed to ascertain statistical significance between different mutant groups.

## Acknowledgements

We are indebted to Prof Mike Lehane for introducing us to the *Drosophila*-trypanosomid parasite work (especially Ed Rowton’s papers) and for his continuous encouragement. We are grateful to Dr Rod Dillon and Dr Lee Haynes for many discussions and technical suggestions that led to various experiments described in this paper. We thank Profs Bruno Lemaitre and Heinrich Jasper for fly stocks. This work was supported by a Consolidator grant from the European Research Council (310912 Droso-Parasite, to P.L.), project grant BB/K003569 from the BBSRC (to P.L.) and a Wellcome Trust doctoral scholarship (to M.A.S).

**Figure SI. Laboratory protocols of *Drosophila* infection with *Herpetomonas muscarum*. (A)** Infection protocol showing the time points for both RNA-seq as well as quantification of parasite numbers following various RNAi treatments. **(B)** Parasite infection was verified also by dissecting guts at different time points. **(C)** Pre-stained parasite with Mitotracker Red was fed to flies that have been fed with DAPI to mark intestinal cells. Stained parasites were seen in the anterior **(C)** and posterior **(D)** midgut. A magnification of D indicated that the parasite (red) were inside the peritrophic matrix (blue) **(E)** A representative standard curve that was made each time with the parasite culture used to infect, so as to help quantify parasite numbers in infection experiments.

**Figure S2. Testing the conditions for RNAi with GAL80^ts^.** Using RFP to investigate the control of GAL80^ts^ over the GAL4 drivers used in this study namely, the general driver J6-GAL4 **(A)**, the EC-specific driver NP1-GAL4 **(B),** the ISC/EB-specific Esg-GAL4 **(C). (D)** At the restrictive temperature (18°C) the system was not inducible following infection.

